# Sequential oxidation of L-lysine by a non-heme iron hydroxylase

**DOI:** 10.1101/2025.01.27.635104

**Authors:** Elizabeth S. Reynolds, Thomas G. Smith, Anoop R. Damodaran, Ambika Bhagi-Damodaran

## Abstract

2-oxoglutarate-dependent non-heme iron hydroxylases offer a direct route to functionalizing C(*sp*^*3*^)–H bonds across a diverse range of substrates, making them prime candidates for chemoenzymatic synthetic strategies. We demonstrate the ability of a non-heme iron L-lysine dioxygenase to perform sequential oxidation and computationally explore structural elements that promote this reactivity.

---

Direct oxidation of C(*sp*^*3*^)–H bonds is often challenging to achieve in a regio- and stereoselective manner as synthetic catalysts frequently struggle to selectively target aliphatic positions over more reactive functional groups.^1^ 2-oxoglutarate-dependent non-heme iron (NHFe) enzymes represent a promising alternative for efficient late-stage C(*sp*^*3*^)– H functionalization of multi-functional molecules, like amino acids, without the need for protecting groups.^2–4^ As hydroxylated amino acids are often used as building blocks for pharmaceutically relevant biomolecules, considerable effort has gone into identifying and engineering NHFe hydroxylases that accept amino acids as their substrates.^2,5–8^

Currently, several NHFe enzymes have been found to hydroxylate free L-lysine at the C3-, C4-, and C5-positions, with the majority of identified species targeting the C4-carbon (**Fig. 1A**).^5,6,9–12^ To better understand the relationship among these previously identified L-lysine 4-hydroxylases, we generated a composite sequence similarity network which reveals three main enzyme populations occupying very distinct sequence space with little overlap even at low sequence identity cutoffs (**Fig. 1B, Fig. S1**). The largest sequence population is defined by KDO2-5 and K4H, (green dots in **Fig. 1B**) hydroxylases which were discovered using genomic mining strategies aimed at identifying members of the clavaminate synthase enzyme superfamily that could hydroxylate amino acids.^6,9,11^ The next sequence population (dark blue dots in **Fig. 1B**) is made up of sequences similar to GlbB, an enzyme responsible for the production of 4*S*-OH-L-lysine in the glidobactin biosynthetic gene cluster.^5^ Finally, the third population (light blue dots in **Fig. 1B**) is composed of sequences similar to lysine dioxygenase (LDO), a NHFe hydroxylase identified based on sequence similarity to an L-lysine-4*R*-halogenase, BesD.^10,13^ While the KDO family members display significant overlap in related sequences, the other two L-lysine-4-hydroxylase populations, defined by GlbB and LDO, exhibit almost no sequence overlap with each other or with the KDOs/K4H, highlighting the distinctiveness of these three populations.

**Fig. 1.**
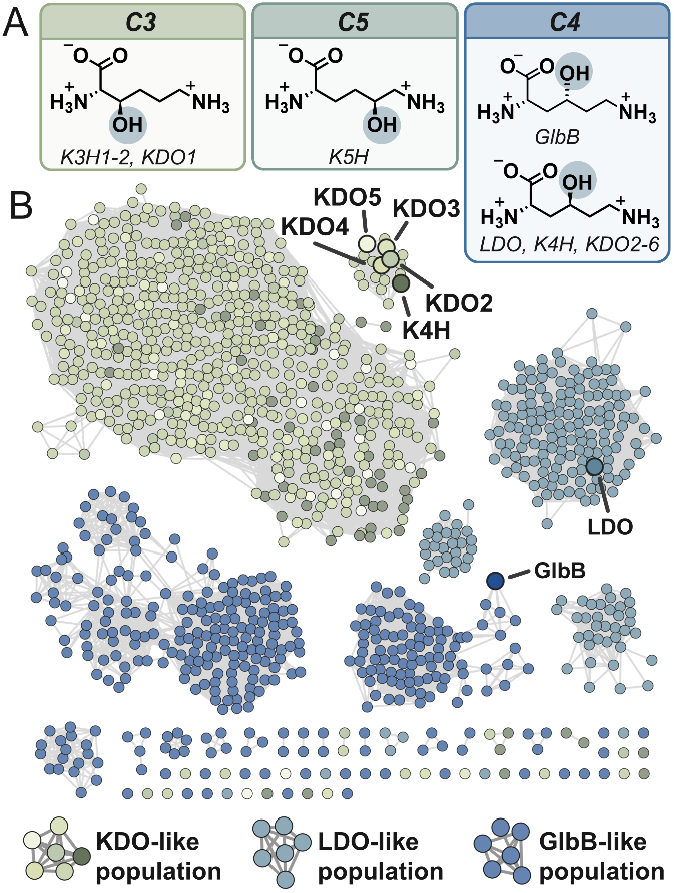
**(A)** L-lysine modifications catalysed by known L-lysine hydroxylases. **(B)** Sequence similarity network constructed from sequences generated from NCBI BLAST search using the known L-lysine-4-hydroxylases as initial queries. Sequence nodes are coloured by which known L-lysine-4-hydroxylase was used as the query for the BLAST search. An alignment score of 40 was used to generate the network which corresponds to ∼35% sequence similarity. Initial query L-lysine-4-hydroxylases are enlarged and labelled.

Here, we focus on LDO (referred to as Hydrox in previous work) and describe its ability to perform multiple oxidations on the native substrate, L-lysine, to generate 4-oxo-L-lysine.^10,14^ Sequential oxidation has been documented in many examples of NHFe enzymes resulting in a variety of products such as aldehydes^15–20^, ketones^21^, carboxylates^15–20^, vicinal diols^22,23^, ether bridges^24^, epoxides^25^, and heterocycles^26^. However, it is not well understood how NHFe enzymes control the extent of substrate oxidation, especially at conditions needed for chemoenzymatic synthesis to be viable. To better understand how LDO enables sequential oxidation at the same carbon to form the oxo-product, we performed Molecular Dynamics (MD) simulations with L-lysine and 4-OH-L-lysine present in the active site. From these, we observed that the addition of the OH-group minimally perturbs the overall substrate orientation, leading to the remaining C4-hydrogen being well positioned for a second abstraction. Overall, using combined biochemical, spectroscopic, and computational strategies, this work characterizes the biocatalytic potential of LDO and explores the structural underpinnings of its sequential oxidation activity.

For initial investigations with LDO, we surveyed its reactivity with L-lysine at various enzyme concentrations and investigated the products through High Performance Liquid Chromatography (HPLC) and mass spectrometry (MS). At a high substrate-to-enzyme ratio (1000:1), we observed ∼50% conversion of L-lysine (**Fig. 2A**, peak shaded blue) to a product with a mass +16 amu relative to the starting material L-lysine (**Fig. 2A**, peak shaded light blue), consistent with the formation of 4-OH-L-lysine as previously characterized (**Table S1**).^10^

**Fig. 2.**
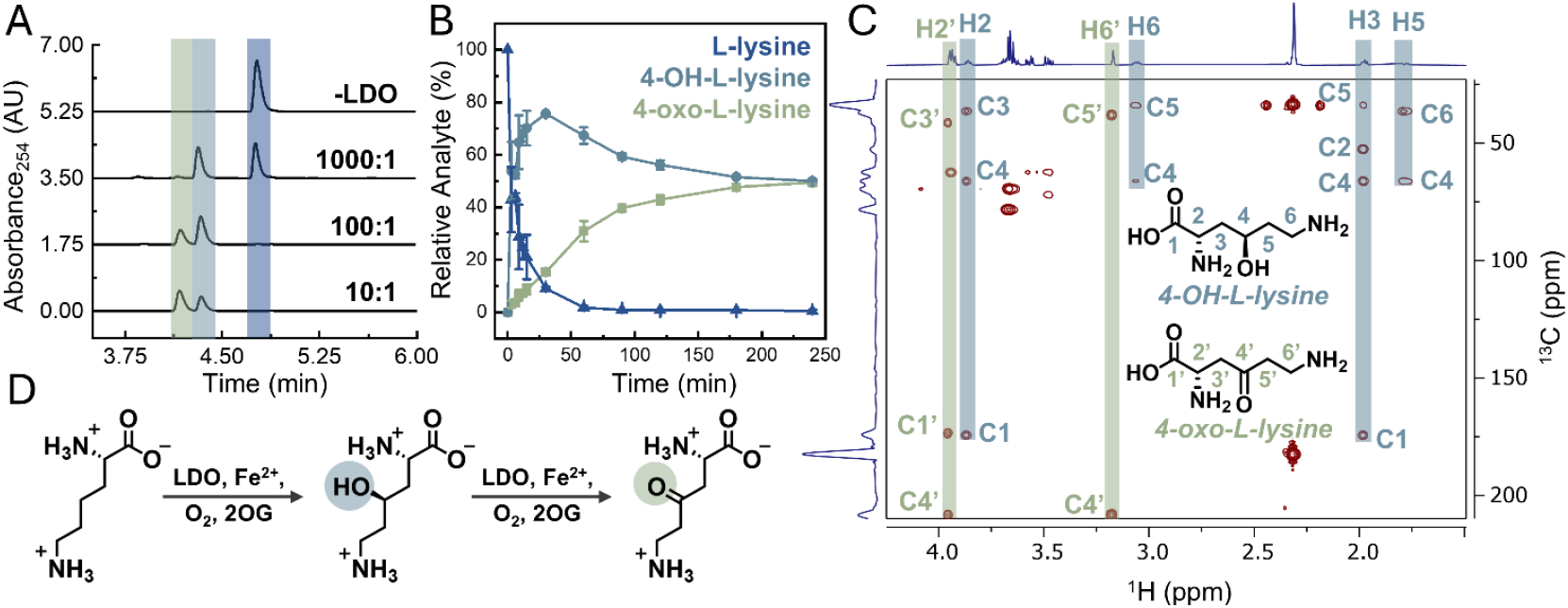
**(A)** HPLC-PDA traces constructed from absorbance at 254 nm for LDO reactions with L-lysine at different substrate-to-enzyme ratios (1000:1, 100:1, 10:1) with peaks corresponding to L-lysine, 4-OH-L-lysine, and 4-oxo-L-lysine highlighted in blue, light blue, and light green, respectively. Identity of products were confirmed by MS from collected HPLC fractions (**Table S1**). **(B)** Time-resolved conversion of L-lysine to 4-OH-L-lysine and 4-oxo-L-lysine by LDO quenched with 0.2 mM EDTA solution. Products detected by HPLC-PDA. **(C)** ^1^H-^13^C HMBC of the enzymatic reaction after 1.5 hours with LDO removed (500 MHz, D_2_O). Highlighted signals have been labeled as corresponding to 4-OH-L-lysine (light blue) and 4-oxo-L-lysine (light green). Only signals for H2’ and H6’ positions are observed for oxo-species as hydrogens at C3’ and C5’ have exchanged with deuterium. Signals that are not highlighted correspond to signals from starting material, succinate, or residual glycerol. **(D)** Proposed sequential oxidation strategy utilized by LDO.

However, as we increased the amount of enzyme relative to substrate (100:1), a secondary product (**Fig. 2A**, peak shaded light green, 41±1% of total product) was formed in addition to mono-hydroxylated species (**Table S1**). An additional increase in the substrate-to-enzyme ratio (10:1), led to the yield of the secondary product (62±1%) surpassing that of 4-OH-L-lysine (38±1%). To better understand the formation of these two products over time, we quenched small volumes of a 50:1 substrate-to-enzyme reaction mixture with an EDTA-containing solution at selected time intervals (**Fig. 2B**). Over the course of the reaction, we observed an initial rapid increase in the concentration of 4-OH-L-lysine, followed by a gradual decline. In contrast, the concentration of the secondary product steadily increased over time before tapering off as 4-OH-L-lysine is seemingly consumed to form this secondary product.

An accurate mass measurement of the new secondary product revealed a mass increase of +14 amu relative to the starting material, L-lysine, which is consistent with either an oxo- or epoxide-product. As both products are known to be formed by NHFe enzymes, we conducted 1D and 2D Nuclear Magnetic Resonance (NMR) characterization (^1^H-^13^C HMBC and HSQC) of the enzymatic reaction to determine the identity of the secondary product.^7,16–20,25^ Within the reaction mixture, we could readily identify ^1^H and ^13^C NMR signals that align with previously reported values for 4*R*-OH-L-lysine (**Fig. 2C**, shaded light blue and **Fig. S2-3**).^10^ Furthermore, we observed NMR signals and reactivity patterns consistent with the assignment of secondary product as 4-oxo-L-lysine (**Fig. 2C**, shaded light green). Most notably, in the ^1^H-^13^C HMBC spectrum of the post-reaction mixture, the ^13^C signal at 208.1 ppm confirmed the presence of a new ketone/aldehyde in the products. Additionally, we observed moderately fast hydrogen-deuterium exchange of several methylene protons of 4-oxo-L-lysine in D_2_O which can be attributed to the increased acidity of the ketone α-hydrogens. While this exchange eliminates some of the proton signals when the reaction is performed in D_2_O, the process of exchange can be confirmed and observed when the reaction is run in buffered H_2_O, lyophilized, and then resuspended in D_2_O immediately before the NMR measurements (**Fig. S4**). From this, we could observe the disappearance of the putative C3’ and C5’ proton signals, as well as the transformation of the C6’ proton signal from a triplet to a singlet over time. The combination of these NMR studies with the results from the enzymatic reaction assays confirm the ability of LDO to sequentially oxidize L-lysine to 4-oxo-L-lysine at moderate to high substrate-to-enzyme ratios.

Recently, KDO3, another L-lysine hydroxylase, was shown to produce a sequentially oxidized L-lysine product.^21^ Despite catalysing nearly identical native reactions, LDO and KDO3 differ significantly in both sequence and structure which suggests that this reactivity is more common than initially characterized in NHFe hydroxylases and is not a unique feature of LDO.^21^ While some examples of sequential oxidation are important for native reactivity^15–18,21–26^, recent attempts to utilize NHFe enzymes for chemoenzymatic synthesis have sometimes struggled to selectively control the extent of oxidation.^7,8,21^ However, it has been demonstrated that the oxidized product outcomes can be manipulated through protein engineering strategies for this class of enzymes. Specifically, as part of their exhaustive 15-round directed evolution campaign of FoPip4H, Cheung-Lee et. al. identified 7 mutations that significantly reduced sequential substrate oxidation, but the exact mechanism by which these mutations improved reaction selectivity was unclear.^7^ A better understanding of how an enzyme allows sequential oxidation of a substrate could lead to more targeted evolution strategies and better support industrial adoption of this class of enzymes. To explore the mechanistic basis for sequential oxidation by LDO, we turned to MD simulations. Like other 2OG-dependent hydroxylases, we anticipate that LDO utilizes a high-valent ferryl intermediate to abstract the H-atom and initiate C–H hydroxylation (**Fig. S5**). To understand how LDO facilitates a second H-atom abstraction, we simulated both L-lysine and 4*R*-OH-L-lysine in the presence of different ferryl intermediate models. From a crystallographic standpoint, LDO is primed to form an off-line ferryl intermediate but reorientation of this intermediate over the course of the reaction has been postulated.^27^ Additionally, while the binding mode of succinate in the ferryl intermediate of TauD has been identified as monodentate, other coordination patterns are thought to be energetically accessible in NHFe hydroxylases.^28,29^ As the exact conformation of the reactive intermediate in LDO has not been characterized, we modeled both in-line and off-line ferryl intermediates with bidentate and monodentate succinate conformations for a total of four different possible ferryl intermediates.

Over the course of the simulations, regardless of ferryl intermediate identity, we found that the orientation of the substrate is impacted only minimally by the presence of the OH-group on 4*R*-OH-L-lysine. Strong interactions formed between the backbone carboxylate of the substrate and surrounding residues H139, W244, and R80 (**Fig. 3A-B**), as the carboxylate engaged in hydrogen bonds with at least one of these residues in 95-100% of the simulations, regardless of substrate identity. Similarly at the opposite end of the molecule, the ε-amine group maintained strong interactions with a well-positioned trio of amino acids, D144, S227, and E125, which engaged in hydrogen bonds with a frequency of 91-99%, 61-93%, and 92-100%, respectively, across all simulations. Finally, more moderate hydrogen bonding patterns are observed with the α-amine, as hydrogen bonds form between residues W143 and D145 for 79-94% and 0-36% of simulations, respectively. Overall, these strong interactions hold L-lysine and 4*R*-OH-L-lysine rigidly in the active site and are not significantly disrupted by the presence of a hydroxyl group at the C4-position (**Fig. 3A-B**). This lack of disruption likely allows for facile binding of both substrates and presents the opportunity for a second proton abstraction to take place.

**Fig. 3.**
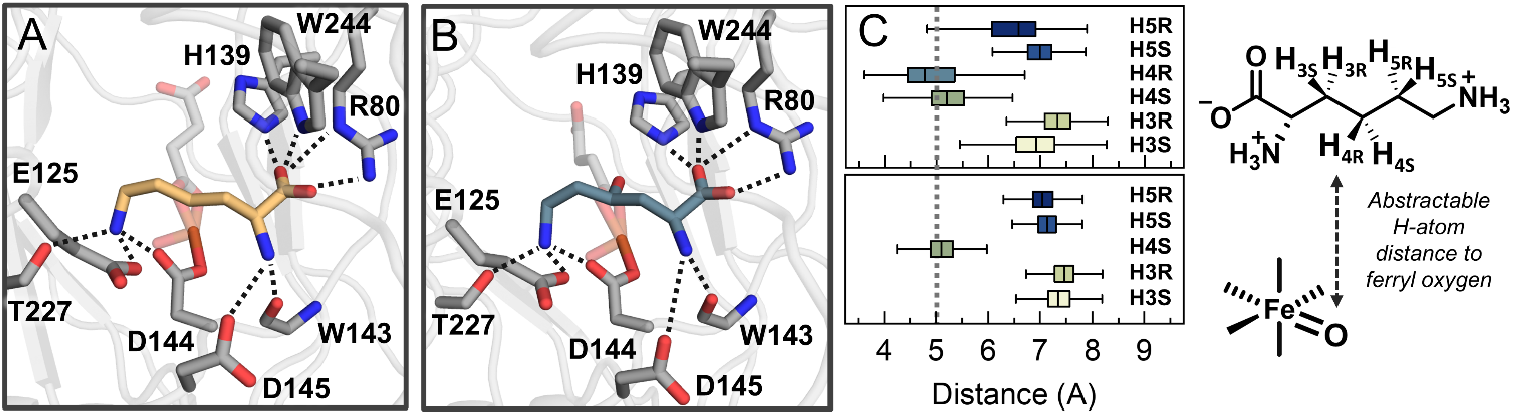
Representative structures from MD simulations with off-line bidentate succinate ferryl intermediate with either **(A)** L-lysine or **(B**) 4*R*-OH-L-lysine, visualizing interactions between the substrate and surrounding active site residues. **(C)** Boxplot of distances from the protons attached to C3-5 carbons of L-Lysine (top) or 4*R*-OH-L-Lysine (bottom) to the oxo-group of the off-line ferryl intermediate with mid-line representing the median. Dashed line is placed at 5 Å which is approximately the threshold for facile H-atom abstraction.

While additional factors that influence reactivity are known to exist, distance to the oxo-group of the ferryl intermediate is typically thought to be a major determinant of which hydrogen is preferentially abstracted.^30–32^ In the simulations with L-lysine, the *pro-R* C4-hydrogen is on average the closest abstractable hydrogen to this oxo-group, and a substrate radical formed at this position agrees experimentally with the observed stereochemistry of the product (**Fig. 3C**). This distance trend holds across all four models of investigated ferryl intermediates (**Fig. S6**). Once the hydroxyl group is present on the substrate, the *pro-S* C4-hydrogen is best poised for abstraction, leading to successive hydroxylation at the same carbon and ultimately the experimentally observed ketone formation. Alternatively, other multi-hydroxylated products or epoxides formed via a mono-hydroxylated intermediate must be able to readily abstract hydrogens at neighbouring carbons (**Fig. S7**).^25^ The hydrogens at the C3- and C5-positions, however, remain farther back on average than the second hydrogen at the C4-position, providing insight as to why sequential oxidation by LDO generates a ketone over an epoxide or a vicinal di-hydroxylated product.

Sequential oxidation has been characterized in a variety of NHFe enzymes, but the factors that allow NHFe hydroxylases to limit or enable sequential oxidation of a substrate are not well understood. As many industrial applications of enzymes require working under very different conditions than those typically found in nature, a deeper understanding of possible product outcomes and the enzymatic mechanisms that enforce these product outcomes are needed. To that end, we describe the sequential oxidation of L-lysine by LDO and examine how the enzyme possibly promotes sequential hydroxylation at the same carbon. From MD simulations, we observed strong interactions specifically with the α-carboxylate and the ε-amine group which enforce very similar orientations on L-lysine and 4*R*-OH-L-lysine, promoting sequential oxidation through close C4-hydrogen positioning. Disruption or promotion of this undifferentiated binding mode of the mono-hydroxylated substrate in future engineering campaigns may enable better control over the degree of product oxidation, allowing for more reliable incorporation of this class of enzymes into biosynthetic pathways.

## Supporting information

SI_sequential_oxidation

## Acknowledgement

ESR acknowledges the support of the National Institute of Health Chemical Biology Training Grant (T32GM132029). Expression, purification and studies with LDO were supported by UMN Startup funds to ABD. Bioinformatics, computational, NMR, biochemical, and mass spec methods in this work were developed via the support of NSF CBET and CLP (Grant # 2046527). Mass spectrometry analysis was performed at The University of Minnesota Department of Chemistry Mass Spectrometry Laboratory (MSL), supported by the Office of the Vice President of Research, CSE, and the Department of Chemistry at the University of Minnesota.

## Conflicts of interest

There are no conflicts to declare.

## Data availability

The data supporting this article have been included as part of the SI.†

## Footnotes

† Electronic supplementary information (ESI) available.

## Notes

### Competing Interest Statement

The authors have declared no competing interest.

