## Supplementary material for "Sequential oxidation of L-lysine by a non-heme iron hydroxylase": SI_sequential_oxidation

### Table of Contents

**Materials and methods**……………………………………………………………………………...S3
Commercial materials………………………………………………………………………………...S3
Protein expression……………………………………………………………………………………...S3
Protein purification……………………………………………………………………………………...S3
Construction of sequence similarity network……………………………………………………...S4
LDO activity assays and product derivatization……………………………………………………...S5
Time-based LDO reaction assay……………………………………………………………………...S5
HPLC detection of reaction products……………………………………………………………...S5
Nuclear magnetic resonance sample preparation and data collection……………………………...S5
Molecular Dynamics Simulations……………………………………………………………………...S6
**Supplementary Tables**……………………………………………………………………………...S7
**Table S1.** Accurate Mass Determination of LDO enzymatic products……………………………....S7
**Supplementary Figures**……………………………………………………………………………...S8
**Figure S1.** Structural comparison of L-lysine-4-Hydroxylases……………………………………S8
**Figure S2.** $^{1}H$-$^{13}C$-HSQC of starting material and succinate standards…………………………S9
**Figure S3.** NMR characterization of LDO enzymatic products from reaction mixture……………S10
**Figure S4.** Deuterium-hydrogen exchange time-course of 4-oxo-L-lysine α-hydrogens……………S11
**Figure S5.** General consensus mechanism for 2OG-dependent hydroxylases……………………S12
**Figure S6.** Abstractable hydrogen distances for MD simulations…………………………………S13
**Figure S7.** Epoxide formation by non-heme iron enzymes…………………………………………S14
**References**……………………………………………………………………………………………S15

### Materials and Methods

#### *Commercial materials*

Imidazole (ReagentPlus®), 2-mercaptoethanol (BME; BioUltra), Magnesium chloride hexahydrate (BioXtra), Amicon Ultra-centrifugal filter units, Sodium Tetraborate Decahydrate (ACS reagent), Ferrous ammonium sulfate hexahydrate (FAS; BioUltra), Ammonium acetate (HPLC grade – LiChropur®), 2-oxoglutarate (2OG; >99.0%), Sodium ascorbate (BioXtra), Acetonitrile (HPLC grade), and Sodium phosphate dibasic (ReagentPlus®) were purchased from Millipore Sigma. BL21-Gold (DE3) and BL21 (DE3) cells were purchased from Agilent and Thermo Scientific, respectively. 6-Aminoquinolyl-N-hydroxysuccinimidyl Carbamate (AQC; >95%) was purchased from Cayman Chemicals. Kanamycin (USP grade), Isopropyl β-D-1-thiogalactopyranoside (IPTG, >99%), RNase A, and Dithiothreitol (DTT; >99.0%) were purchased from GoldBio. Tryptone (BioReagent), Yeast extract (BioReagent), Calcium chloride (ACS reagent), DNase I solution, Sodium chloride (ACS reagent), Ethylenediaminetetraacetic acid disodium salt dihydrate (EDTA; BioReagent), L-lysine monohydrochloride (BioReagent), and Tris Hydrochloride (Molecular Biology) were purchased from Fisher Scientific. HEPES (>99.5%) was purchased from DOT Scientific. $D_2O$ (99.9%) was purchased from Cambridge Isotope Laboratories. Reagent grade or purity is given in parentheses if applicable. HisTrapFF-5 ml columns were purchased from Cytiva. NMR tubes were purchased from VWR. All HPLC solvents were either HPLC grade or filtered through 0.22 µm filters.

#### *Protein Expression*

A previously constructed plasmid[1] for His10-tagged LDO from *Streptomyces roseifaciens* (referred to as Hydrox in previous work[2]) was used for subsequent studies. A plasmid containing a His-tagged human rhinovirus 3C protease (also commercially known as PreScission™ Protease when fused with glutathione S-transferase) with N-terminal domain of the spider silk protein construct (NT*-HRV3CP) was obtained from Addgene.[3] Plasmids containing LDO and NT*-HRV3CP were transformed into BL21-Gold (DE3) and BL21 (DE3) competent cells, respectively. For expression of LDO, colonies of transformed cells from an overnight LB-agar plate ([kanamycin] = 50 µg/mL) were inoculated into primary 2XYT cultures (50 mL, [kanamycin] = 50 µg/mL) and incubated at 37 °C overnight at 200 RPM. Culture flasks with 2XYT media (1.5 L, [kanamycin] = 50 µg/mL) were inoculated with overnight cultures and grown at 37 °C at 200 RPM until cells reached an OD 600 of 0.6−0.8. After reaching suitable optical densities, protein expression was induced with IPTG (0.5 mM), and then cultures incubated at 18 °C for 24 hours. Expression for NT*-HRV3CP was performed following similar methodology to LDO expression except TB media was used instead of 2XYT and secondary cultures were incubated for 20 hours after induction with IPTG. Cells were collected by centrifugation at 6000 RPM at 4 °C, flash-frozen, and stored at −20 °C for further use.

#### *Protein purification*

For purification of LDO, frozen cells were resuspended in lysis buffer (300 mM NaCl, 10 mM imidazole, 10% (vol/vol) glycerol, 5 mM $CaCl_2$, 5 mM $MgCl_2$, 0.1 mg/mL RNase, and 0.05 U/mL DNase 1, 50 mM HEPES (pH=7.5)) and then lysed by sonication. The lysed cells were centrifuged at 20,000 RPM at 4 °C, and the resulting supernatant was filtered through 0.45 µm filters. Filtered supernatant was applied to a 5mL HisTrapFF column using an AKTA Start protein purification system. After sample application, the column was washed with 10 column volumes (CVs) of wash buffer (300 mM NaCl, 10 mM imidazole, 10% (vol/vol) glycerol, 20 mM 2-mercaptoethanol, 50mM HEPES (pH= 7.5)), and then the His-tagged LDO was eluted using a two-step isocratic method first with 28% elution buffer (300mM NaCl, 300mM imidazole, 10% glycerol (vol/vol), 20 mM 2-mercaptoethanol, 50mM HEPES (pH 7.5)) for 8 CVs followed by 8 CVs of 100% elution buffer. Fractions from the second elution phase were collected and dialyzed in 100 mM NaCl, 1 mM DTT, 1 mM EDTA, 50 mM HEPES (pH =7.5) for 2 hours, followed by another 2-hour dialysis in 1 mM DTT, 100 mM NaCl, 50 mM HEPES (pH =7.5). After this second round of dialysis, the protein solution was diluted to a concentration of 0.02−0.05 mM and then incubated with NT*-HRV3CP protease (1 mg protease to 100 mg protein), dialyzing against 1 mM DTT, 100 mM NaCl, 50 mM HEPES (pH = 7.5) overnight. The dialyzed protein solution was then concentrated to approximately 10 mL using an Amicon Ultra-15 centrifugal filter (10 kDa MWCO). Imidazole was added to the protein solution at a final concentration of 10 mM, and then protein solution was applied to a 5 mL HisTrapFF column using an AKTA Start protein purification system. His-tagged cleaved LDO was recovered during sample application and addition 4 CVs of wash buffer application. The collected solution was buffer exchanged into 50 mM HEPES (pH = 7.5) through 5 washes in an Amicon Ultra-15 centrifugal filter (10 kDa MWCO). After the final wash, the protein solution was concentrated, and glycerol was added to the protein solution for a final concentration of 10% (vol/vol). After concentration of the protein was assessed at 280 nm using extinction coefficient calculated by ExPASy[4], the protein was aliquoted, flash-frozen, and stored at −80 °C for further use.

For NT*-HRV3CP purification, frozen cells were resuspended in lysis buffer (300 mM NaCl, 10 mM imidazole, 5% (vol/vol) glycerol, 5 mM $CaCl_2$, 5 mM $MgCl_2$, 0.1 mg/mL RNase, and 0.05 U/mL DNase1, 50 mM Tris-HCl (pH 7.5)). Cells were lysed, centrifuged, filtered, and applied to 5 mL HisTrapFF column as described for LDO. After sample application, the column was washed with 10 column volumes (CVs) of wash buffer (300 mM NaCl, 10 mM imidazole, 5% (vol/vol) glycerol, 50 mM Tris-HCl (pH 7.5)), and then the His-tagged NT*-HRV3CP was eluted using a two-step isocratic method with first 10% elution buffer (300 mM NaCl, 500 mM imidazole, 5% (vol/vol) glycerol, 50 mM Tris-HCl (pH 7.5)) for 3 CVs followed by 8 CVs of 100% elution buffer. Fractions from the second elution phase were collected and dialyzed in 300 mM NaCl, 1 mM DTT, 50 mM Tris-HCl (pH 7.5) for a total of three rounds of two-hour dialysis. The solution from dialysis was concentrated using an Amicon Ultra-15 centrifugal filter (10 kDa MWCO). After concentration, glycerol was added to the protein solution for a final concentration of 10% (vol/vol) and concentration was assessed 280 nm using extinction coefficient calculated by ExPASy.[4] The protein was then aliquoted, flash-frozen, and stored at −80 °C for further use.

***Construction of sequence similarity network***

Sequences for LDO[2] (WP_107105619), GlbB[5] (AKJ29081.1), KDO2[6] (ACU60313.1), KDO3[6] (ABQ06186.1), KDO4[6] (AEV99100.1), KDO5[6] (J3BZS6), and K4H-4 (EFK34737.1) were used as inputs for NCBI BLAST search against the nonredundant protein sequence database with an expect threshold of 0.05 and max target sequences set to 5000. Resulting similar sequences for all Lysine-4-hydroxylases were combined into a single set, and sequences with 65% similarity were removed by CD-HIT.[7] Using this condensed sequence list as input, a sequence similarity network (SSN) was constructed using the EFI-EST web tool with an alignment score cutoff of 40, which corresponded to a sequence identity of ~35%, a common starting threshold for separating isofunctional homologous enzymes.[8,9] The SSN was visualized with Cytoscape v.3.10.2.[10]

#### *LDO activity assays and product derivatization*

Before activity assays, LDO was buffer exchanged into 50 mM HEPES (pH 7.5) using 0.5 mL Amicon™ Ultra 10 kDa MWCO centrifugal filter units at 13500 RPM at 4˚ C for 15-minute wash cycles to remove storage glycerol. For varied substrate-to-enzyme ratio studies, reaction samples consisted of 5 mM L-lysine, 600 μM FAS, 20 mM 2OG, 5 mM sodium ascorbate with either 5, 50, or 500 μM LDO. All reaction samples (50 μl) were incubated in an Eppendorf Thermomixer® at 25˚ C for 5 hours while mixing (450 RPM).

After incubation, reactions were diluted 10-fold in 50 mM HEPES (pH 7.5) and then were filtered through 0.5 mL Amicon™ Ultra 10 kDa MWCO centrifugal filter units at 13500 RPM for 20 minutes at 4˚ C. Reaction sample flowthrough (80 μL) was diluted 2-fold in 15 mM sodium tetraborate (pH 10.5). Two sequential additions (20 μL) of AQC in acetonitrile (3 mg/ml) were added to each reaction (final volume: 200 μL), and samples were vortexed to combine after each addition. Derivatized samples were then analyzed immediately or stored at -80 or -20 ˚ C for future analysis.

#### *Time-based LDO reaction assay*

LDO was buffer exchanged in 50 mM HEPES (pH 7.5) as described above. A reaction (350 μl) was prepared with the following conditions: 2.5 mM L-lysine, 300 μM FAS, 20 mM 2OG, 5 mM sodium ascorbate, and 50 μM LDO in 50 mM HEPES (pH 7.5). Aliquots (20 μL) of reaction sample were quenched by 10x dilution in 200 μM EDTA in 50 mM HEPES (pH 7.5) solution at indicated time points. Reaction aliquots (200 μL) were filtered to remove LDO and derivatized with AQC as described in previous section.

#### *HPLC detection of reaction products*

Derivatized amino acid samples were analyzed using a Shimadzu Prominence-i LC-2030C 3D Plus system equipped with a Poroshell 120 CS-C18 column (3.0 mm x 150 mm x 2.7 μm) and a PDA detector. For all samples, the column temperature was 35°C, the flow rate was set to 0.6 mL/min, and sample injection volume was 5 μL. Solvent A was 5 mM ammonium acetate (pH = 5.0) and Solvent B was 60% acetonitrile in water. Gradient conditions were: 20% B, 0.0 to 1.5 min; 20% B to 75% B, 1.5 to 5.5 min; 75% B to 100% B, 5.5 to 6.0 min; 100% B, 6.0 to 7.0 min; 100% B to 20% B, 7.0 to 7.5 min; 20% B, 7.5 to 10.0 min. All HPLC traces were constructed from absorbance at 254 nm. Peak area integrations were performed in Shimadzu Lab Solution software. Percent analyte conversion was determined by the peak area of the analyte divided by the sum of all analytes peak areas. To determine the identity of reaction products, samples of HPLC chromatographic peaks were collected and diluted in methanol. Mass spectra of analytes were obtained by direct infusion using a syringe pump onto a Sciex X500R QTOF-MS system with electrospray ionization in positive ionization mode.

#### **Nuclear magnetic resonance sample preparation and data collection**

All reactions were incubated in an Eppendorf Thermomixer® at 25˚ C for the indicated amount of time while mixing at 450 RPM. For structural characterization of 4-OH-L-Lysine and 4-oxo-L-Lysine studies,

LDO was buffer exchanged into 50 mM sodium phosphate in $D_2O$ (pD 7.5) using 0.5 mL Amicon™ Ultra (10 kDa MWCO) centrifugal filter units at 13500 RPM at 4˚ C for five 15-minute wash cycles to remove storage glycerol. The reaction was prepared using stock solutions of compounds in 50 mM sodium phosphate in $D_2O$ (pD 7.5) so that final reaction conditions were 5 mM L-lysine, 300 μM FAS, 10 mM 2OG, 10 mM sodium ascorbate with 200 μM LDO in 50 mM sodium phosphate in $D_2O$ (pD 7.5). After 1.5-hour incubation, the reaction sample was filtered through 0.5 mL Amicon™ Ultra (10 kDa MWCO) centrifugal filter units at 13500 RPM for 20 minutes at 4˚ C to remove LDO. Flowthrough was transferred to 5 mm O.D. NMR tubes. Starting material and succinate standards were made from diluted stock solutions used to make the reaction sample. For hydrogen-deuterium exchange studies, all stock solutions were prepared in 50 mM sodium phosphate in $H_2O$ (pH 7.5), and a reaction sample was prepared with final concentrations of 5 mM L-lysine, 300 μM FAS, 15 mM 2-oxoglutrate, 10 mM sodium ascorbate with 200 μM LDO in 50 mM Sodium phosphate in $D_2O$ (pD 7.5). After a 1.5-hour incubation at 25 °C, the sample was filtered to remove LDO as described previously, and then the sample was lyophilized to remove $H_2O$. Sample was resuspended in $D_2O$ immediately prior to initial spectra collection to observe hydrogen-deuterium exchange at α-hydrogens of putative 4-oxo-L-lysine. 1H, 1H-13C HSQC, and 1H-13C HMBC NMR spectra were acquired at 25 °C on a Bruker 500 MHz Avance III HD spectrometer equipped with a 5 mm Prodigy TCI-H&F Cryoprobe.

***Molecular Dynamics Simulations***

The starting structure of LDO was taken from the protein data bank (PDB: 7JSD; Chain C).[2] Missing residues were added using the MODELLER module[11] in UCSF ChimeraX[12] and hydrogen atoms were added to the protein using a pH of 7.5 with the H++ Web server.[13–15] The crystal structure active site was modified using PyMOL[16] and Avogadro[17] to generate the four ferryl intermediate models. LDO structures with 4*R*-OH-L-lysine were made by modifying the bound crystallized L-lysine structure *in-silico* using Avogadro[17]. Force field parameters describing covalent bonds to ferryl iron for each of these models were generated with the MCPB.py module.[18,19] Protein residues in LDO were parameterized with the AMBER ff19SB force field.[20,21] Succinate and substrates were treated with the generalized amber force field (GAFF) with missing parameters developed using the Antechamber package.[22,23] The protein was solvated in a 12.0 Å OPC water octahedron cell, and counterions $Na^+$ were added to a concentration of 0.15 M, followed by the addition of $Cl^-$ to neutralize the system.[24] Protein was minimized using a seven-step minimization process where solute heavy atoms were subject to restraints starting at 10.0 kcal/mol/Å$^2$, which were decreased geometrically until reaching 0.0 kcal/mol/Å$^2$. The system was then gently heated to 300 K and density equilibrated for 3.5 ns at a constant temperature of 300 K. After equilibration, three unrestrained independent 100 ns simulations were performed for each ferryl intermediate and substrate combination. CPPTRAJ[25] was used to perform all trajectory analyses with H-bond donor/acceptor distance cutoff set to 3.2 Å.

**Supplementary Tables**

**Table S1.** Accurate Mass Determination of LDO enzymatic products

| Analyte | HPLC retention time (min) | Chemical formula | Theoretical $[M+H]^+$ | Observed $[M+H]^+$ | Mass error (ppm) |
|---|---|---|---|---|---|
| AQC(2x)-OH-L-lysine | 4.17 | $C_{26}H_{26}N_6O_5$ | 503.2037 | 503.2069 | 6.36 |
| AQC(2x)-oxo-L-lysine | 4.34 | $C_{26}H_{24}N_6O_5$ | 501.1881 | 501.1886 | 1.00 |

**Supplementary Figures**

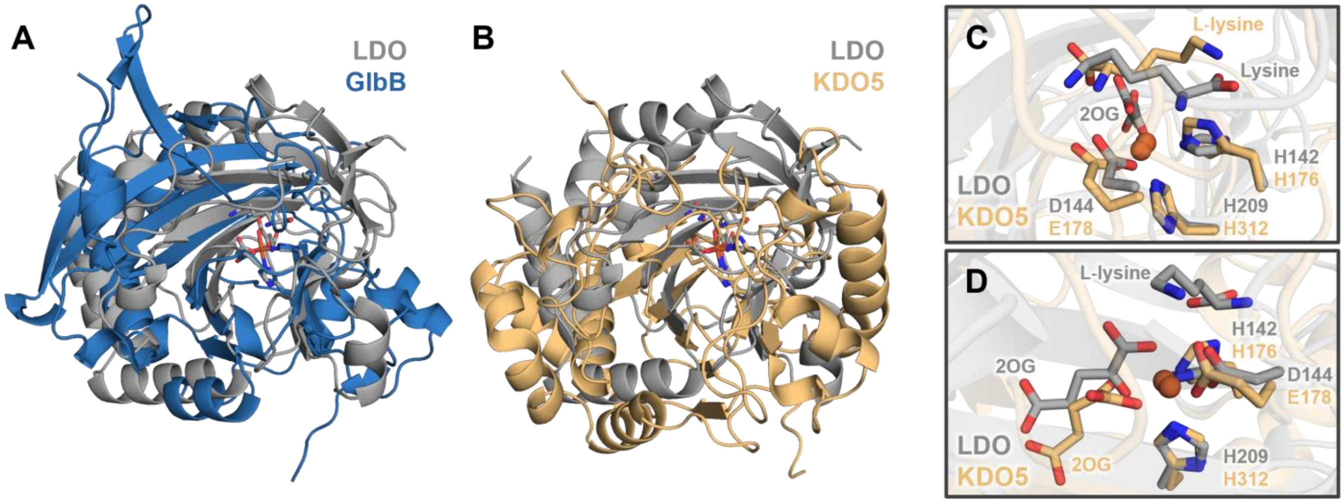


**Figure S1.** Structural comparison of L-lysine-4-hydroxylases. **(A)** Alphafold[26] structure of GlbB overlaid with crystal structure of LDO (7JSD - chain a) aligned by residues of the facial triad (GlbB: H188, D190, and H251; LDO: H142, D144, and H209) due to lack of structural similarity. **(B)** Crystal structure of KDO5 (6EXF - chain B) overlaid with crystal structure of LDO (7JSD - chain A) aligned by histidines of the facial triad and crystallized Fe (KDO5: H176 and H312; LDO: H142 and H209) due to lack of structural similarity. **(C)** Overlaid active site structures of KDO5 (6EXF - chain B) and LDO (7JSD - chain A) showcasing difference in L-lysine binding orientation **(D)** Overlaid active site structures of KDO5 (6EUR - chain B) and LDO (7JSD - chain A) showcasing differences in 2OG orientation.

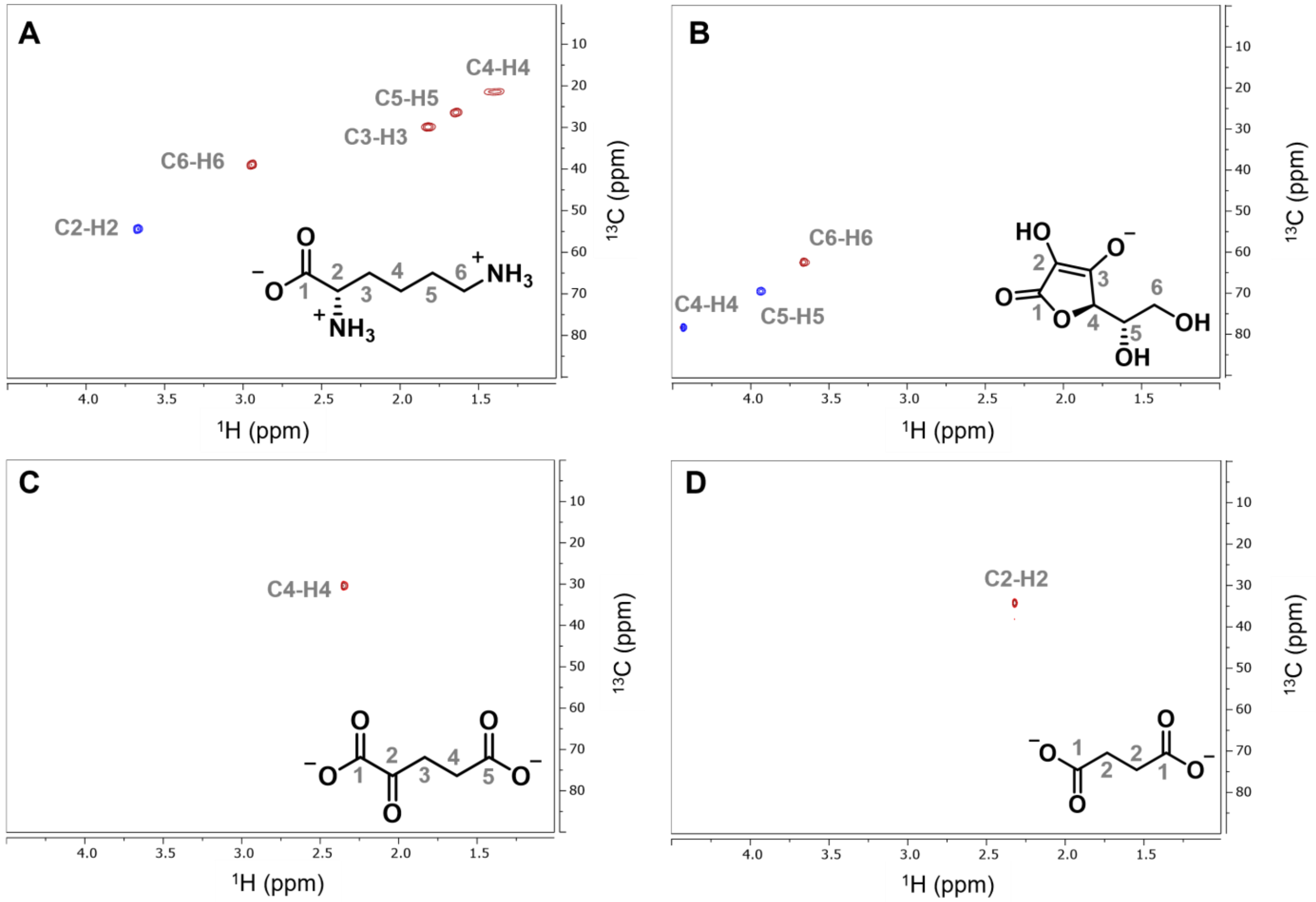


**Figure S2.** $^1H$-$^{13}C$ HSQC of starting material and succinate standards. **(A)** 5 mM L-lysine HCl, **(B)** 10 mM Sodium Ascorbate, **(C)** 10 mM 2-oxoglutaric acid (C3-H3 signal is eliminated due to hydrogen-deuterium exchange of α-hydrogens of ketone), and **(D)** 10 mM Sodium succinate in 50 mM Sodium Phosphate (pD = 7.5) (500.15 MHz, D2O). For HSQC experiments, -CH and -CH3 signals are in-phase (blue) while -CH2 signals are anti-phase (red).

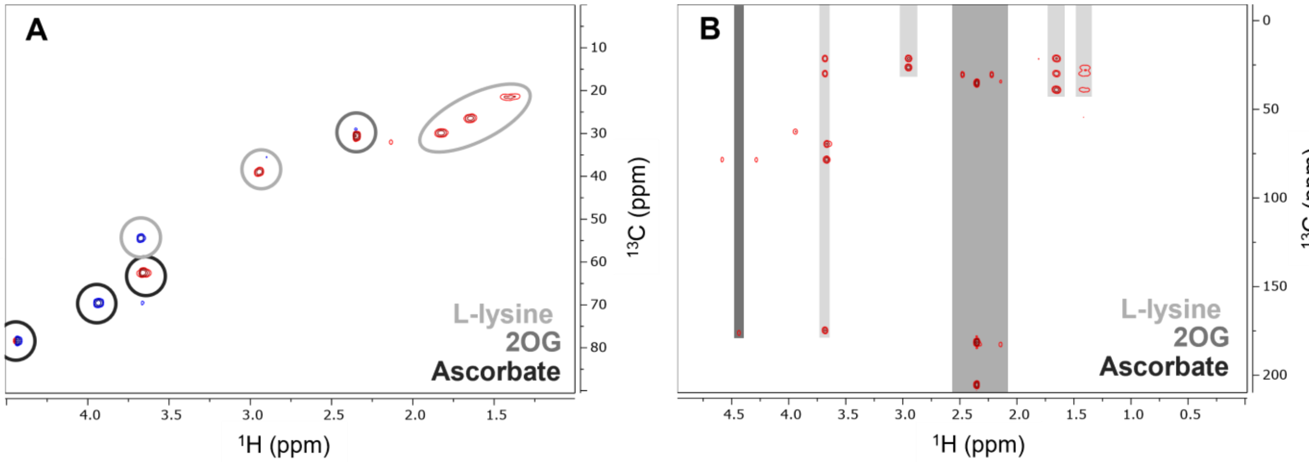


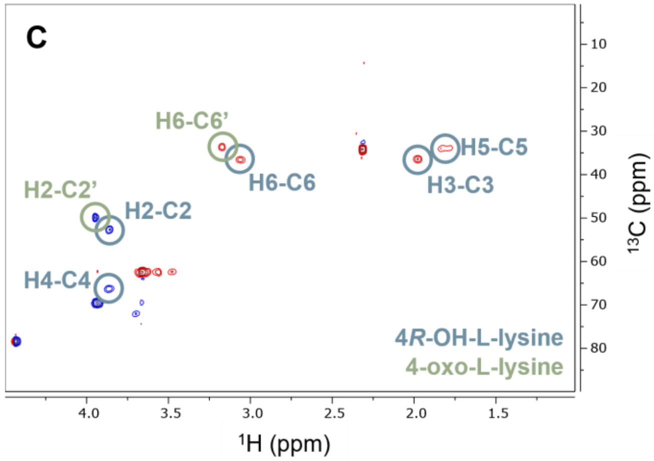


D

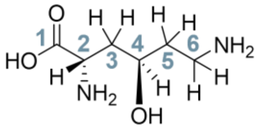

*4R-OH-L-lysine*

| | $^{1}$H (ppm) | $^{13}$C (ppm) |
|---|---|---|
| 1 | --- | 174.2 |
| 2 | 3.86 | 52.6 |
| 3 | 1.98 | 36.3 |
| 4 | 3.86 | 66.3 |
| 5 | 1.80 | 34.1 |
| 6 | 3.06 | 36.5 |

E

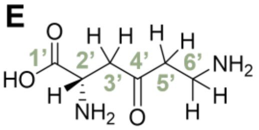

*4-oxo-L-lysine*

| | $^{1}$H (ppm) | $^{13}$C (ppm) |
|---|---|---|
| 1' | --- | 173.4 |
| 2' | 3.96 | 49.9 |
| 3' | 3.14* | 41.2 |
| 4' | --- | 208.1 |
| 5' | 2.96* | 38.2 |
| 6' | 3.17 | 33.6 |

*These proton chemical shifts were observed during a deuterium exchange study, see Figure S4

**Figure S3**. NMR characterization of LDO enzymatic products from reaction mixture. (**A**) $^{1}$H-$^{13}$C HSQC of reaction starting materials without iron (5 mM L-lysine-HCl, 10 mM 2-oxoglutarate, 10 mM Sodium Ascorbate in 50 mM sodium phosphate buffer (pD = 7.5)) (500.15 MHz, D2O) (**B**) $^{1}$H-$^{13}$C HMBC of reaction starting materials without iron (5 mM L-lysine-HCl, 10 mM 2-oxoglutarate, and 10 mM Sodium Ascorbate in 50 mM sodium phosphate buffer (pD = 7.5)) (500.15 MHz, D2O). (**C**) $^{1}$H-$^{13}$C HSQC of enzymatic reaction after 1.5 hours with LDO removed (500.15 MHz, D2O). Reaction conditions: 200 μM LDO, 300 μM FAS, 5 mM L-lysine-HCl, 10 mM 2-oxoglutarate, and 10 mM Sodium Ascorbate in 50 mM Sodium Phosphate buffer (pD = 7.5) in $D_2O$. Chemical shifts determined from the NMR data for (**D**) 4*R*-OH-L-lysine and (**E**) 4-oxo-L-lysine. 4*R*-OH-L-lysine chemical shifts match those determined previously in literature.[2] For HSQC experiments, -CH and -$CH_3$ signals are in-phase (blue) while -$CH_2$ signals are anti-phase (red).

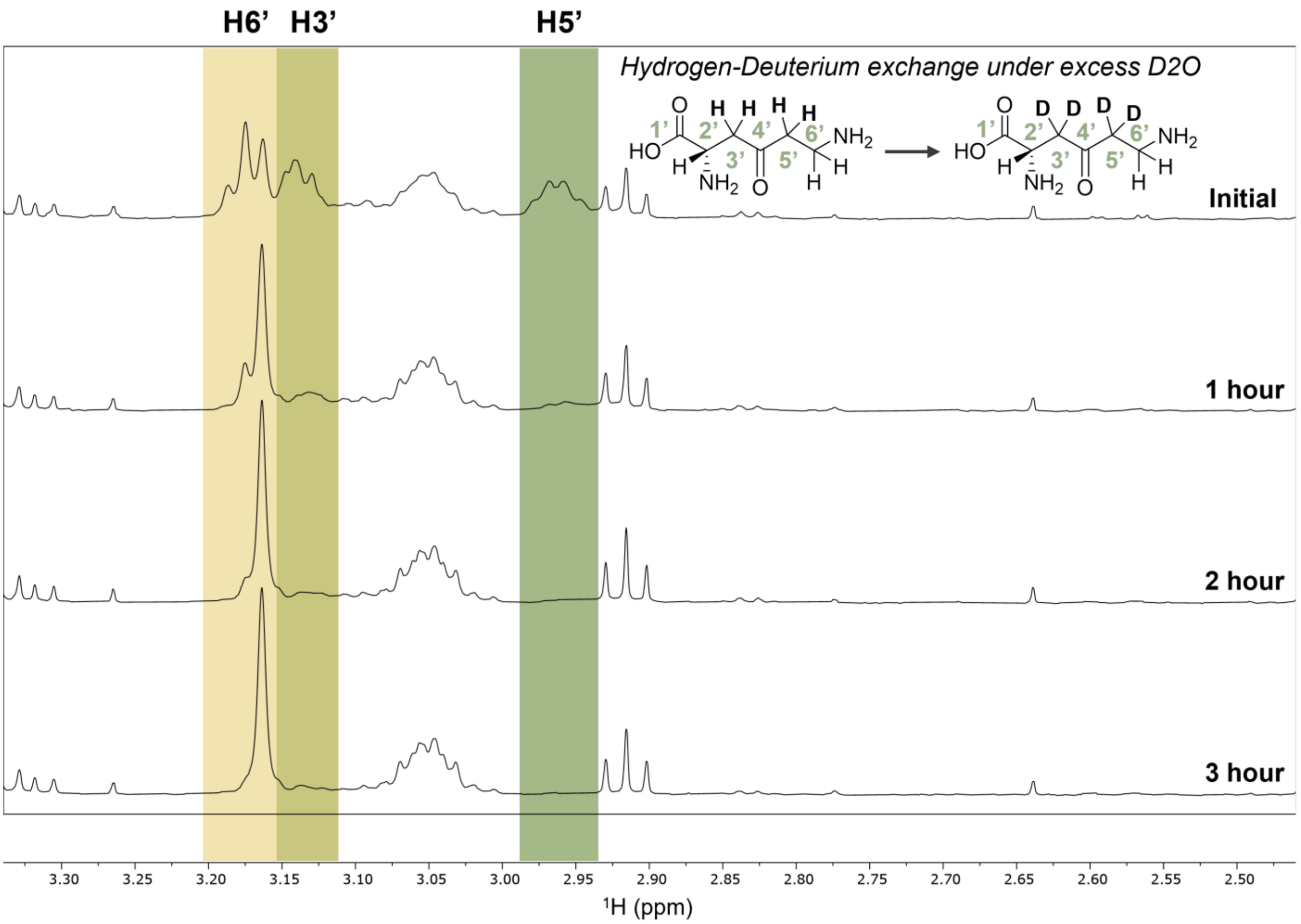


**Figure S4**. Deuterium-hydrogen exchange time-course of 4-oxo-L-lysine α-hydrogens. $^{1}$H-NMR of enzymatic reaction after 1.5 hours with LDO removed (500.15 MHz, $D_2O$). Reaction conditions: 200 μM LDO, 300 μM ferrous ammonium sulphate, 5 mM L-lysine-HCl, 15 mM 2-oxoglutarate, and 10 mM Sodium Ascorbate in 50 mM sodium phosphate buffer (pH = 7.5) in $H_2O$. After reaction, LDO was removed, and sample was lyophilized. Sample was reconstituted in $D_2O$ immediately prior to initial spectra collection and measurements were taken every hour after that point.

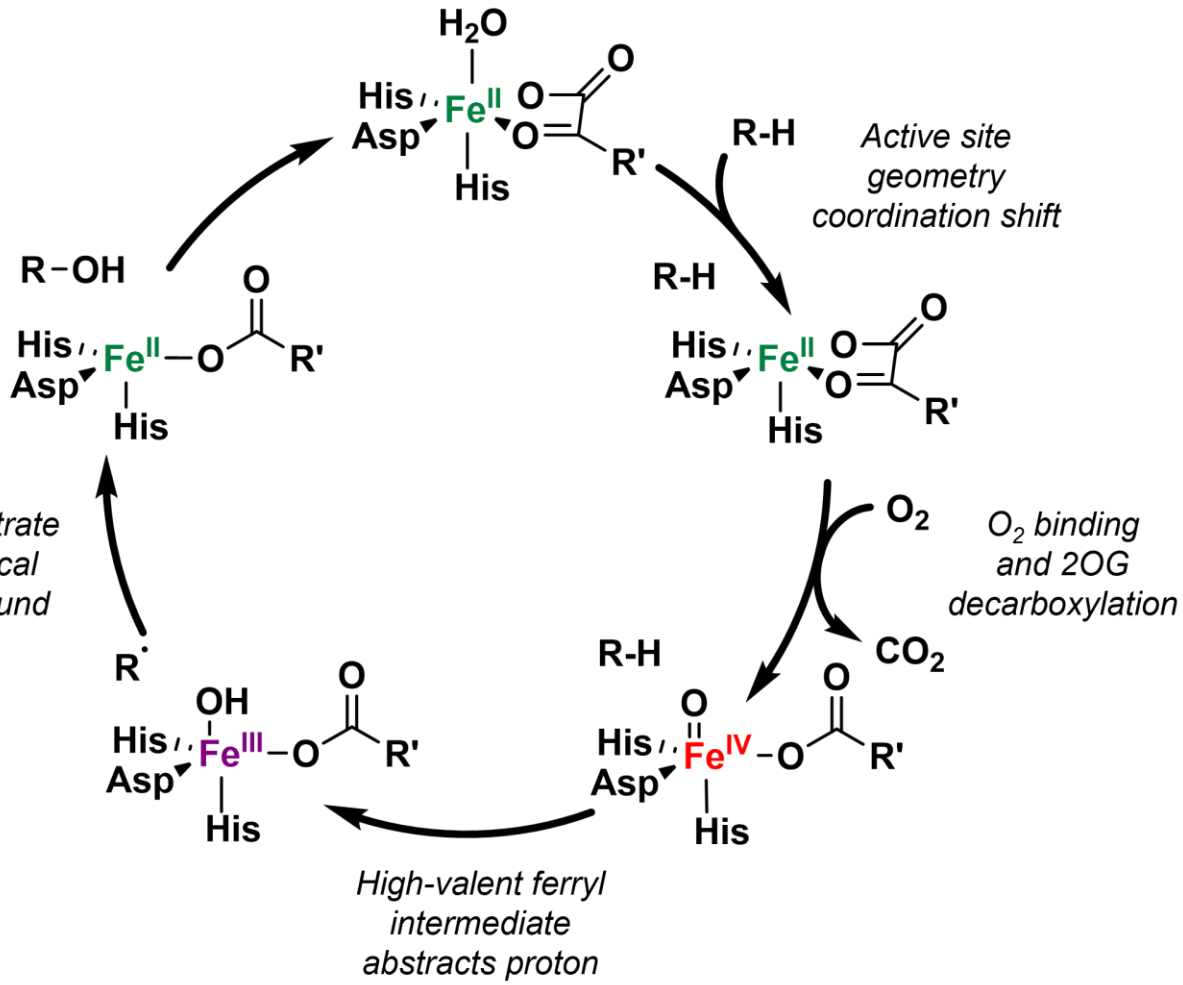


**Figure S5.** General consensus mechanism for 2OG-dependent hydroxylases. In the resting state of the enzyme, the iron is coordinated by an aqua ligand, 2OG, and protein derived histidine and aspartate ligands. Upon substrate binding, the aqua ligand is displaced and the active site coordination shifts from a 6-coordinate environment to a 5-coordinate environment to allow space for efficient oxygen binding. Upon $O_2$ binding, co-substrate 2OG is decarboxylated, enabling the formation of the high valent ferryl intermediate. This highly reactive intermediate then abstracts a hydrogen atom from the substrate, forming a substrate radical which then rebounds with the hydroxyl ligand to form the hydroxylated product.

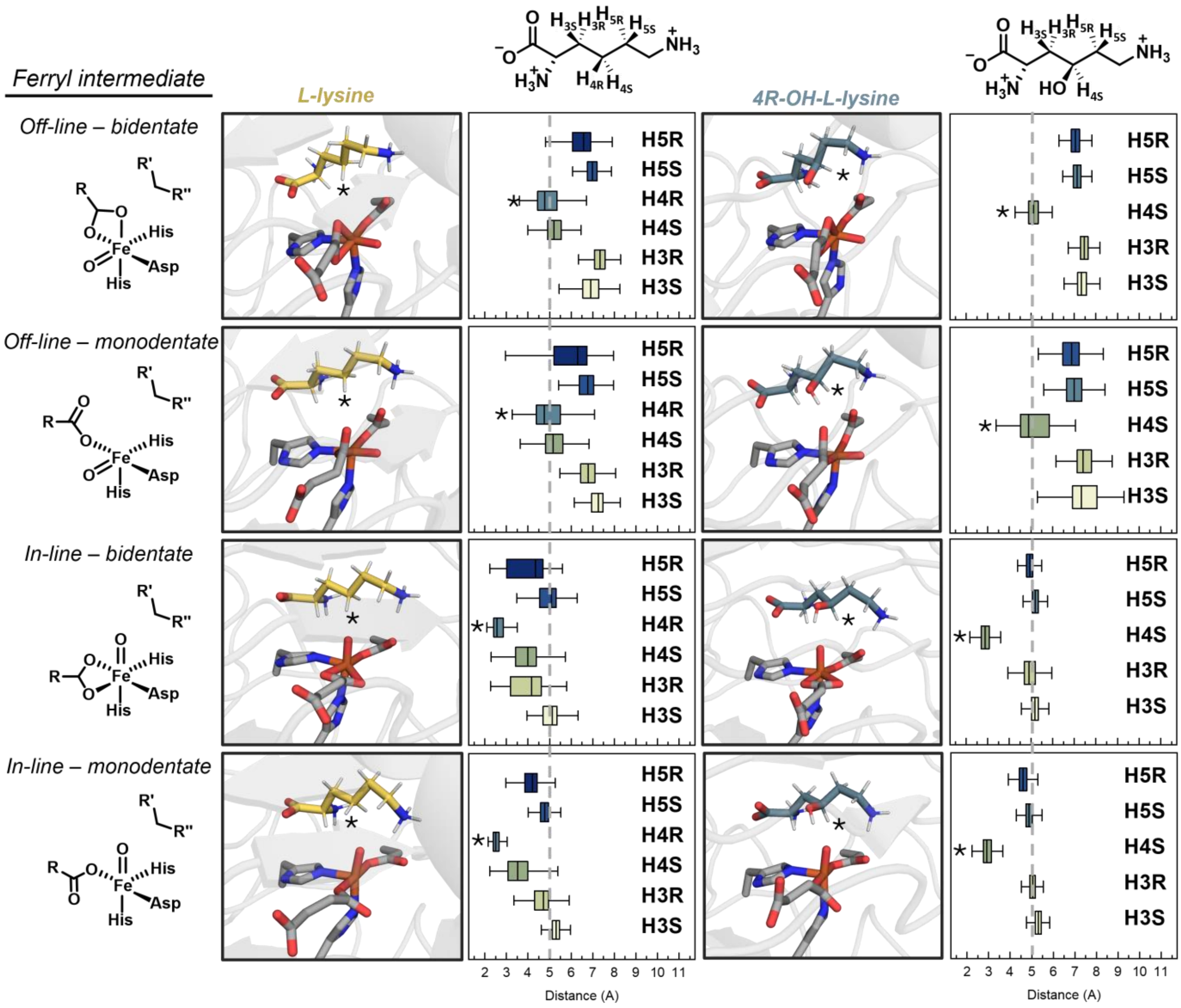


**Figure S6**. Abstractable hydrogen distances for MD simulations. Distance from the protons attached to C3-5 carbons of L-lysine (yellow) or 4*R*-OH-L-lysine (gray blue) to the oxo-group of the ferryl intermediate for the four different models. Closest proton is denoted by an asterisks in the structure and the corresponding boxplot. Dashed line is shown at 5 Å which is approximately the threshold for facile hydrogen atom abstraction.

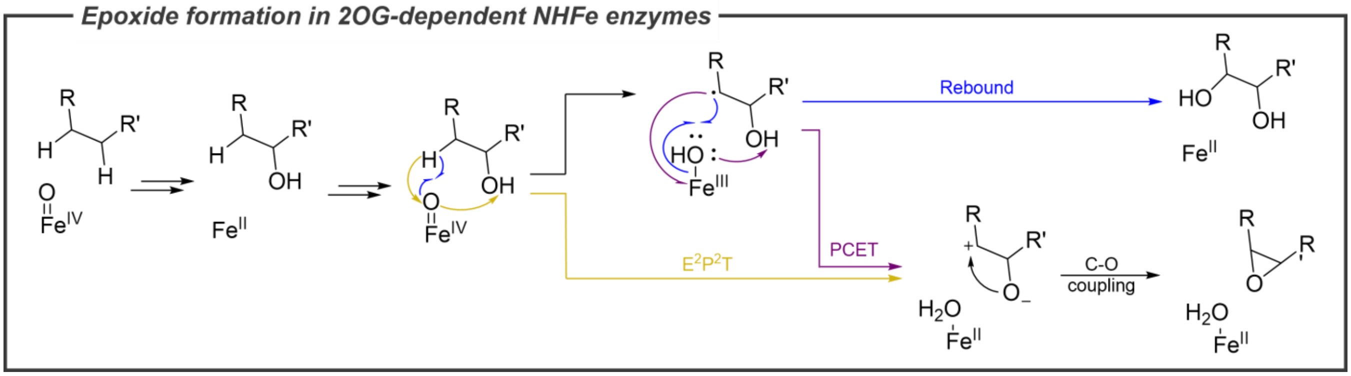


**Figure S7.** Epoxide formation by non-heme iron enzymes through a mono-hydroxylated intermediary product as proposed for Hyoscyamine 6β-hydroxylase (H6H).[27] Other routes to epoxide formation for 2OG-dependent non-heme iron enzymes involve an initial alkene product, as in the case of AsqJ.[28]
